# Reoxygenation after Evofosfamide Treatment in Pancreatic Ductal Adenocarcinoma Xenografts is due to Decreased Oxygen Consumption and not Increased Oxygen Supply

**DOI:** 10.1101/2020.05.16.099853

**Authors:** Shun Kishimoto, Jeffrey R. Brender, Yu Saida, Kazutoshi Yamamoto, James B. Mitchell, Murali C. Krishna

## Abstract

Evofosfamide is designed to release a cytotoxic bromo-isophosphoramide (Br-IPM) moiety in a hypoxic microenvironment. This drug therefore preferentially attacks hypoxic regions in tumors where other standard anti-cancer treatments such as chemotherapy and radiation therapy are often ineffective. Various combination therapies with evofosfamide have been proposed and tested in preclinical and clinical settings. However, the treatment effect of evofosfamide monotherapy on tumor hypoxia has not been fully understood, partly due to the lack of quantitative methods to assess tumor pO2 in vivo. Here, we use quantitative pO2 imaging by EPR to evaluate the change in tumor hypoxia in response to evofosfamide treatment using two pancreatic ductal adenocarcinoma xenograft models; MIA Paca-2 tumors responding to evofosfamide and Su.86.86 tumors which do not respond. EPR imaging showed oxygenation improved globally after evofosfamide treatment in hypoxic MIA Paca-2 tumors, in agreement with the ex vivo results obtained from hypoxia staining by pimonidazole and in apparent contrast to the decrease in K^trans^ observed in DCE MRI. This suggests reoxygenation after treatment is due to decreased oxygen demand rather than improved prefusion. Following the change in pO2 after treatment may therefore yield a way of monitoring treatment response. The observation that evofosfamide not only kills the hypoxic region of the tumor but also improves oxygenation in the residual tumor regions provides a rationale for combination therapies using radiation and anti-proliferatives post evofosfamide for improved outcomes.

## Introduction

Tumors are able to grow to a size of 2-3 mm^3^ by relying on passive diffusion of oxygen and nutrients; for further growth and eventual metastasis additional vasculature is required which is developed by recruiting angiogenic pathways to develop *de novo* vasculature.^1^ While in normal processes there is a tight control between anti-angiogenic and pro-angiogenic pathways which results in a well-structured and functional vascular network, in tumors the pro-angiogenic factors dominate to meet the tumor growth demands. This results in a tumor vascular network which is both poorly organized structurally and functionally abnormal.^1^ As a consequence, the tumor microenvironment is characterized by poor perfusion, low oxygen (hypoxia), and high interstitial fluid pressure.^2,3^ Additionally, tumor cells often develop other altered chemical characteristics such as elevated aerobic glycolysis and low pH.^4^ The eventual development of such a harsh microenvironment results in tumors resistant to treatment with chemotherapy and radiotherapy.^5^ Solid tumors are known to have regions of hypoxia which render them resistant to radiotherapy.^3^ Hypoxia activated prodrugs (HAPs) have been developed recently to target hypoxic regions of the tumor which are normally resistant to radiation and anti-proliferative chemotherapeutic agents.^6^ The hypoxia activated prodrug evofosfamide is comprised of a nitroimidazole moiety covalently linked to a bromo-iphopsharamide (Br-IPM) moiety^7,8^. Intracellularly, the nitroimidazole moiety can undergo one-electron reduction to the corresponding anion radical through cellular reductases. Once formed, the anion radical undergoes rapid reoxygenation at diffusion limited rates in normoxic conditions, returning back to the original state. Under these conditions, the prodrug formulation is retained and evofosfamide does not exert significant cytotoxicity as the reactive Br-IPM moiety is blocked. However, under hypoxic conditions the equilibrium between the reduced and oxidized forms is shifted, and evofosfamide is susceptible to fragmentation in the reduced form to release the pharmacologically active Br-IPM moiety. Once generated, free Br-IPM is a powerful alkylating agent capable of cross-linking DNA and inducing apoptosis. *In vitro*, evofosfamide has been tested in various cell lines and found to have selective hypoxic cytotoxicity.^7,8^ Based on several pre-clinical studies, evofosfamide has been tested clinically in a phase I/phase II study in refractory multiple myeloma as a hypoxia activated prodrug and found to have shown efficacy.^9^

A few pre-clinical studies have explored the use of evofosfamide in combination therapies.^10–15^ When evaluating combination therapies involving oxygen dependent treatments such as radiation therapy, it is crucial to know the pO2 levels during the lifecycle of the treatment as radiation therapy is not well tolerated and should be administered at the point of maximum efficacy. As evofosfamide likely changes these levels, the timing of evofosfamide administration can be manipulated to maximize the treatment efficacy during planning of combination therapy. The evidence for an evofosfamide induced decrease in the hypoxic fraction of the tumor rests on ex vivo immunohistochemical analyses,^16^ which are qualitative and require serial tissue biopsies, which may be difficult to implement clinically. To plan the treatment regimen effectively, a non-invasive method to evaluate the change in pO2 in response to evofosfamide quantitatively would be useful.

EPRI (Electron Paramagnetic Resonance Imaging) is one of the most reliable methods of measuring pO2 *in vivo* in pre-clinical models.^17,18^ EPRI requires the injection of a non-toxic paramagnetic spin probe and absolute pO2 can be calculated from the linewidth of the distributed probe. The paramagnetic spin probe is well tolerated for serial imaging without impacting the physiology or biochemical profile of the tissue being interrogated. With EPRI it is possible to experimentally determine both the fractional volume of the hypoxic region of the tumor with pO2 < 10 mmHg and the pO2 spatial distribution.^19^ It is also possible to dynamically monitor changes in tumor oxygenation by EPRI using hyperoxygenation strategies or by chemical hypoxia induction.^13,18,20^ EPRI has been previously used in preclinical studies to examine the pretreatment tumor oxygenation status quantitatively to examine the efficacy of evofosfamide in combination with pyruvate to induce hypoxia or in combination with radiation.^13,14^ pO2 values from EPRI were successful in predicting evofosfamide sensitivity across cell lines.

These studies showed the importance of determining the pretreatment tumor oxygenation status quantitatively for effective treatment. While evaluating combination therapies that involve oxygen dependent treatment such as radiation therapy, it is also crucial to understand the pO2 change in response to evofosfamide in tumor microenvironment because the residual fraction of the tumor microenvironment can be modified by the treatment. In such case, we need to consider the timing of evofosfamide administration to maximize the treatment efficacy when combination therapy is planned. Recent studies suggested that evofosfamide induced a decrease in hypoxic fraction, such effects were monitored ex vivo using histochemical assays method which are qualitative. ^21^ While planning treatments with radiation or anti-proliferatives after evofosfamide. it is important to investigate the difference in pO2 between pre- and post-treatment with evofosfamide in a unique tumor.

In this study, to evaluate the changes in oxygenation after evofosfamide treatment, tumor blood perfusion was examined using dynamic contrast enhanced MRI (DCE MRI). Combined with the pO2 information obtained by EPRI, we could achieve the comprehensive assessment of treatment efficacy with evofosfamide by non-invasive imaging modalities. The reoxygenation effect after evofosfamide treatment was found to be driven by a decrease in oxygen consumption due to cell death or cell arrest rather than improved prefusion from relief of solid stress on the vascular network.

## Material and Methods

### Chemicals

Evofosfamide was purchased from Threshold Pharmaceuticals. Gd-DTPA was purchased from Bio-PAL, Inc. The triarylmethyl EPR oxygen tracer OX063 (methyl-tris[8-carboxy-2,2,6,6-tetrakis[2-hydroxyethyl]benzo[1,2-d:4,5-d0]-bis[1,3]dithiol-4-yl]-trisodium salt) was obtained from GE Healthcare.

### In vitro cytotoxicity assay

Exponentially growing cells were seeded in a 6 well plate (10000 cells/well) 6 hours before the addition of evofosfamide. After the drug addition, the cells were incubated under aerobic conditions for 48 hours at 37°C in a standard tissue culture incubator. Cellular cytotoxicity was assessed by counting viable cells after treatment and normalized to control conditions.

### Animal experiments

All animal experiments were conducted in compliance with the Guide for the Care and Use of Laboratory Animal Resources (National Research Council, 1996), and the experimental protocols were approved by the National Cancer Institute Animal Care and Use Committee (RBB-159-2SA). Female athymic nude mice were supplied by the Frederick Cancer Research Center, Animal Production. human pancreatic adenocarcinoma MIA Paca-2 cells and Su.86.86 (obtained from Threshold Pharmaceuticals) were authenticated in May 2013 by RADIL using a panel of microsatellite markers. Cells were routinely cultured in RPMI 1640 with 10% FCS. The tumors were formed by injecting 3 × 10^6^ subcutaneously into the right hind legs of female athymic nude mice. Tumor-bearing mice were treated daily with the intra-peritoneal administration of 50 mg/kg evofosfamide five days a week when tumor size reached approximately 400 mm^3^. In the imaging experiments, mice were anesthetized by isoflurane inhalation (4% for inducing and 1%–2% for maintaining anesthesia) and positioned prone with their tumor-bearing legs placed inside the resonator. During EPRI and MRI measurements, the breathing rate of the mouse was monitored with a pressure transducer (SA, Instruments, Inc.) and maintained at 60 +/− 20 breaths/min. Core body temperature was maintained at 36 +/− 1 °C with a flow of warm air.

### Immunohistochemistry

Frozen tumor sections were thawed at room temperature, then fixed with ice cold acetone for 10 minutes. Following blocking, the sections were incubated with Hypoxyprobe^TM^ rabbit anti-pimonidazole antibody (1:250; hpi) for pimonidazole staining or rat anti-mouse CD31 antibody (1:250; BD) for CD31 staining overnight at 4℃. Fluorescence microscopy and imaging was performed using a BZ-9000 BIOREVO (KEYENCE). Images were captured with the BZ-9000E viewer at 10X magnification and were stitched to compose a whole image of the sections using the BZ-II Analyzer. Quantification of the pimonidazole-positive was completed by counting the pixels of the positive area.

### Electron paramagnetic resonance imaging

The technical details of the EPR scanner and oxygen image reconstruction were described in earlier reports.^22–25^ Homemade resonators tuned to 300 MHz were used for EPRI. After the mouse was placed in the resonator, the EPR oxygen tracer OX063 (1.125 mmol/kg bolus) was injected intravenously under isoflurane anesthesia. The repetition time was 8.0 μs. The free induction decay (FID) signals were collected following the radiofrequency excitation pulses under a nested looping of the x, y, and z gradients and each time point in the FID underwent phase modulation enabling 3D spatial encoding. Since FIDs last for 1 to 5 μs, it is possible to generate a sequence of T2* maps that is, EPR linewidth maps, which linearly correlate with the local concentration of oxygen and enable the pixel-wise estimation of pO2.

### Dynamic contrast enhancement MRI

DCE-MRI studies were performed on a 1.0T scanner (ICON; Bruker Bio-Spin MRI GmbH). T1-weighted fast low angle shot (FLASH) images were obtained with TE = 6 ms, TR = 118 ms, a flip angle of 30°, two slices, 0.4 · 0.4 mm^2^ resolution, 15 s acquisition time per image, and 45 repetitions. Gd-DTPA solution (0.25 mmol/kg body weight) was injected through a tail vein cannula 1 min after the start of the dynamic FLASH sequence. To determine the local concentrations of Gd-DTPA, T1 maps were calculated from three sets of Rapid Imaging with Refocused Echoes (RARE) images obtained with TR = 500, 1,000, and 2,000 ms, with the acquisitions being made before running the FLASH sequence. The endothelial transfer coefficient K^trans^ was calculated by fitting the dynamic MRI signal change to the standardized Tofts model.^26,27^

### Statistics

Data were expressed as the means +/− standard deviation. The paired Student’s t-test was used to compare the values before and after treatment in the same group of mice in EPRI and MRI studies. Other comparisons of means, which required independent sets of mice, were analyzed using the independent Student’s t-test. The log rank test was used to compare the distribution of Kaplan-Meier survival curves. p < 0.05 was considered statistically significant.

## Results

The differential treatment response to evofosfamide was examined using two human PDAC xenografts that were previously indicated to be sensitive (MIA Paca-2) and resistant (Su.86.86) to evofosfamide therapy.^7,8^ Su.86.86 cells stained higher for CD31, a marker for blood vessels, than the MIAPaca-2 cell line (Figure 1A, B) consistent with prior studies which indicated MIA Paca-2 xenografts are more hypoxic in vivo compared to corresponding Su.86.86 xenografts.^14,19^ We confirmed the sensitivity of each cell line by observing tumor growth after the initiation of the evofosfamide treatment (Fig. 2A and Fig. 2B). A statistically significant reduction in tumor growth compared with the groups treated with vehicle (5% DMSO in PBS) is evident in the Kaplan-Meier plots in the MIA Paca-2 tumors treated with 50 mg/kg evofosfamide five days a week (Fig. 2C: p=0.02, log rank test) but no observable inhibition in Su.86.86 was noticed (Fig. 2D: p=0.8). These results were also mirrored in *in vitro* studies under aerobic conditions (Figure 3). Both cell types were effectively killed at micromolar concentrations of evofosfamide in ambient air equilibrated condition, suggesting residual activity still exists even under high oxygen concentrations. Furthermore MIA Paca-2 cells were still killed at lower concentrations of evofosfamide than Su.86.86 cells, even in the *in vitro* situation in which differences in oxygen delivery are not expected to be a limiting factor, although consumptive hypoxia cannot be ruled out *a priori*.^28^

**Figure 1.**
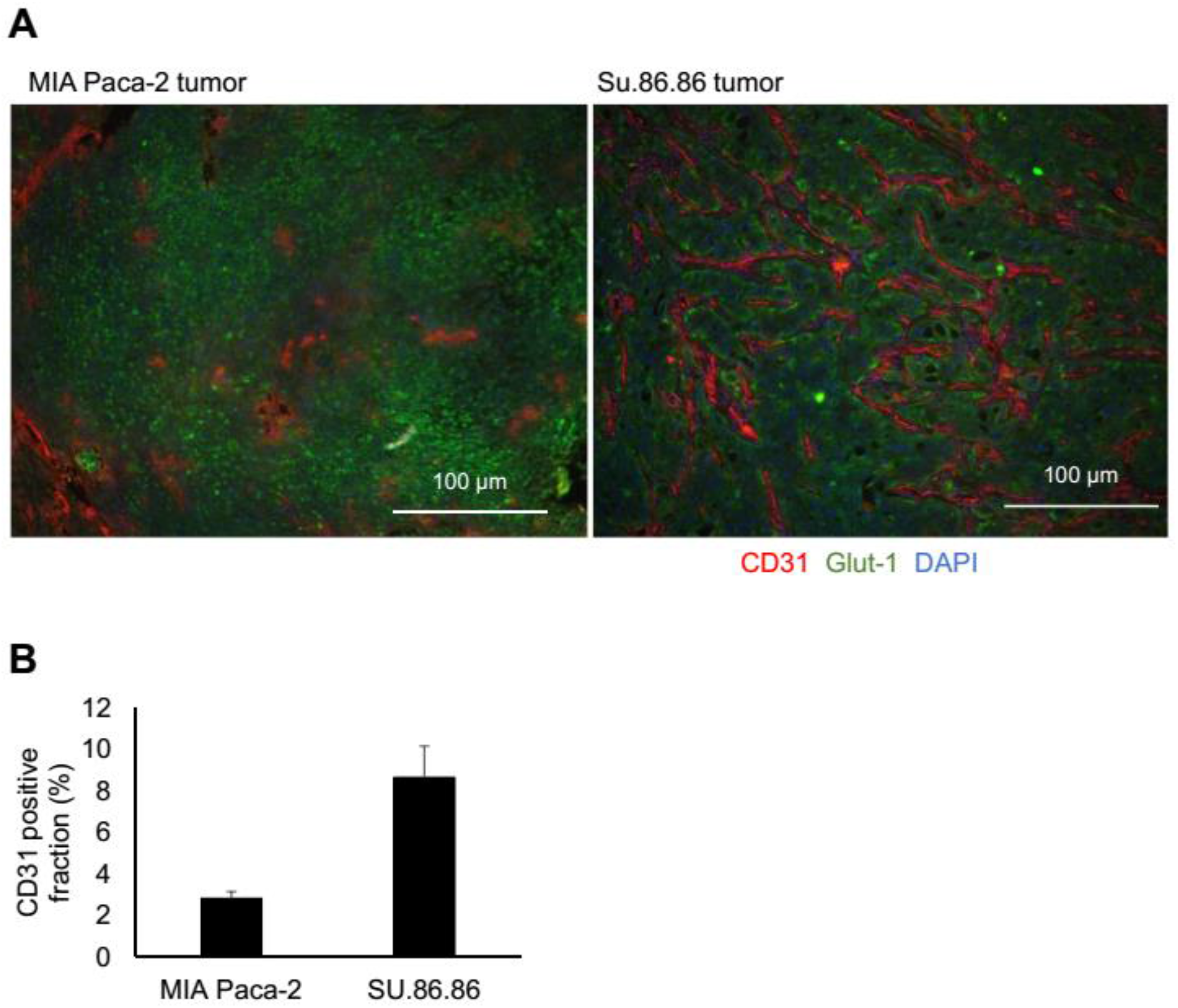
Difference in vascularity between poorly differentiated MIA Paca-2 tumor and highly differentiated Su.86.86 tumor evaluated by immunohistochemistry assays. **A.** CD31, Glut-1, and DAPI was stained in red, green, and blue, respectively. **B.** CD31 positive fraction for both tumors. The fraction was calculated from randomly selected 5 fields per tumor section (n = 4 per group).

**Figure 2.**
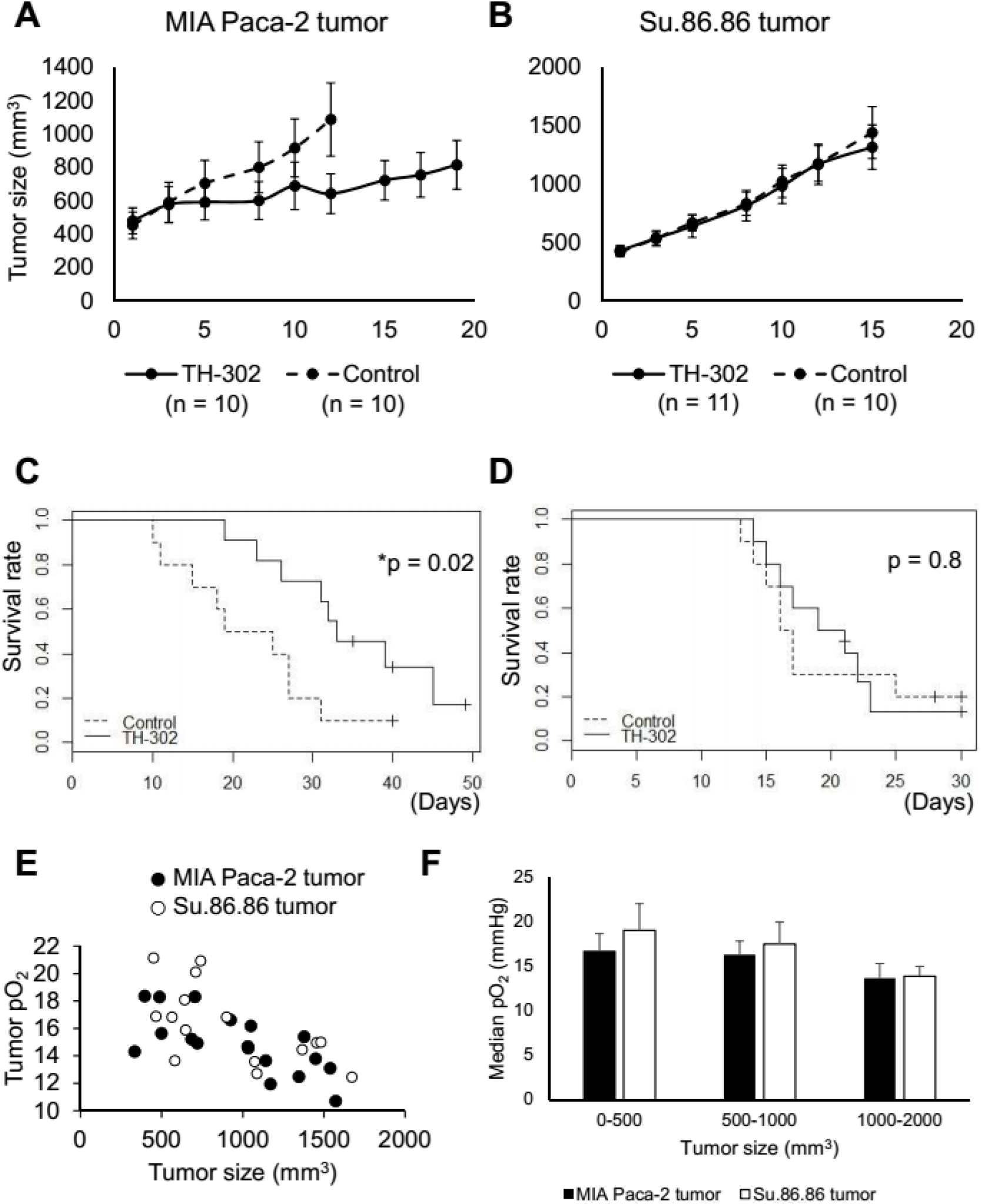
Assessment of evofosfamide treatment efficacy on MIA Paca-2 and Su.86.86 cells and tumors. **A, B** The tumor growth curve of MIA Paca-2 and Su.86.86 tumors treated with evofosfamide (evofosfamide 50 mg/kg i.p. 5 days/week) or vehicle. The number of animals is indicated in the plot. **C, D** Kaplan-Meier survival curves of Fig.2A, B experiments, respectively. * shows the p<0.05 evaluated by the log-rank test. **E.** The relationship between tumor size and tumor pO2 for both MIA Paca-2 and Su.86.86 tumors evaluated by EPR oximetry.

**Figure 3.**
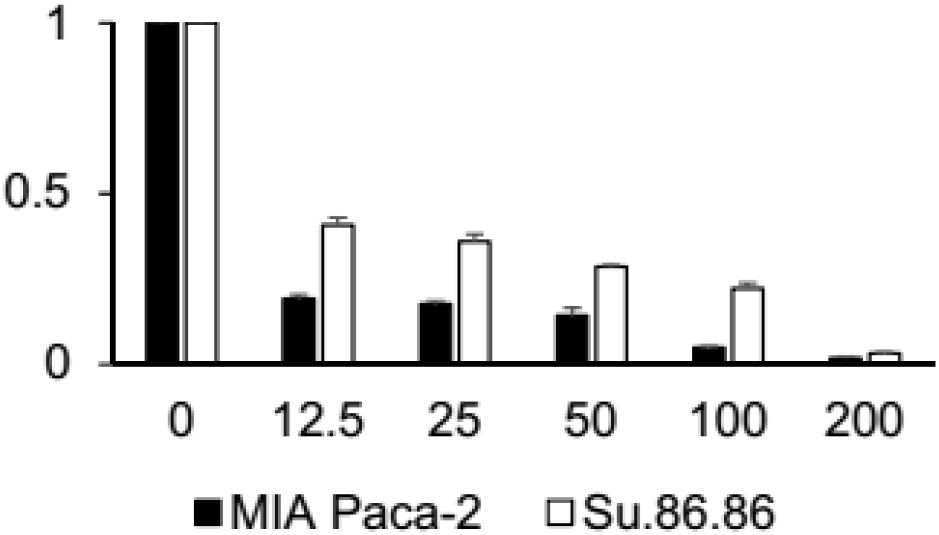
In vitro cytotoxicity evaluated after 48-hour of incubation with evofosfamide at the indicated concentrations (0, 12.5, 25, 50, 100, 200 μM).

The proposed mechanism for evofosfamide cytotoxicity involves a reduction of the inactive pro-drug to generate the reactive species, which is reversed under normoxic conditions.^7,8^ Since the equilibrium of the fragmentation process generating the pharmacologically active species from the inactive prodrug is dependent on the local oxygen levels, the efficacy should be correlated with the hypoxic fraction. To assess relative hypoxia in the two xenograft models, EPR imaging studies of these tumors were performed. Earlier studies have shown that Su.86.86 had lower hypoxic fractions than MIA Paca-2 tumors, consistent with the higher staining for CD31 in Su.86.86 (Figure 1B), a marker for blood vessels, a trend noticed in current experiments as well. As expected, there was a correlation between tumor size and oxygen levels (Figure 2E and F). Considering the oxygen dependent activation of evofosfamide, this difference in oxygenation status prior to treatment may lead to a difference in treatment response. This assessment assumes the hypoxia is unchanged with treatment. However, the cytotoxic effect of evofosfamide is expected to lower oxygen consumption while apoptotic or necrotic cell death is expected to relieve the interstitial pressure within the tumor, increasing oxygen supply by alleviating the constriction of the capillaries supplying blood to the tumor.^3^

To determine the functional alteration in the tumor microenvironment induced by evofosfamide in MIA Paca-2 tumors, we examined tumor hypoxia using pimonidazole staining on evofosfamide treated and untreated tissue sections. Substantial differences in pimonidazole positive area between untreated and evofosfamide treated tumors were found, in agreement with previous study evaluating tumor hypoxia after evofosfamide treatment (Figure 4A).^21^ In general, it is accepted that pimonidazole binds tissue at the threshold of pO_2_ < 10 mmHg.^16^ Using this threshold, the pimonidazole stained area decreased by more than 50% in evofosfamide treated tumors (Fig. 4B).

**Figure 4.**
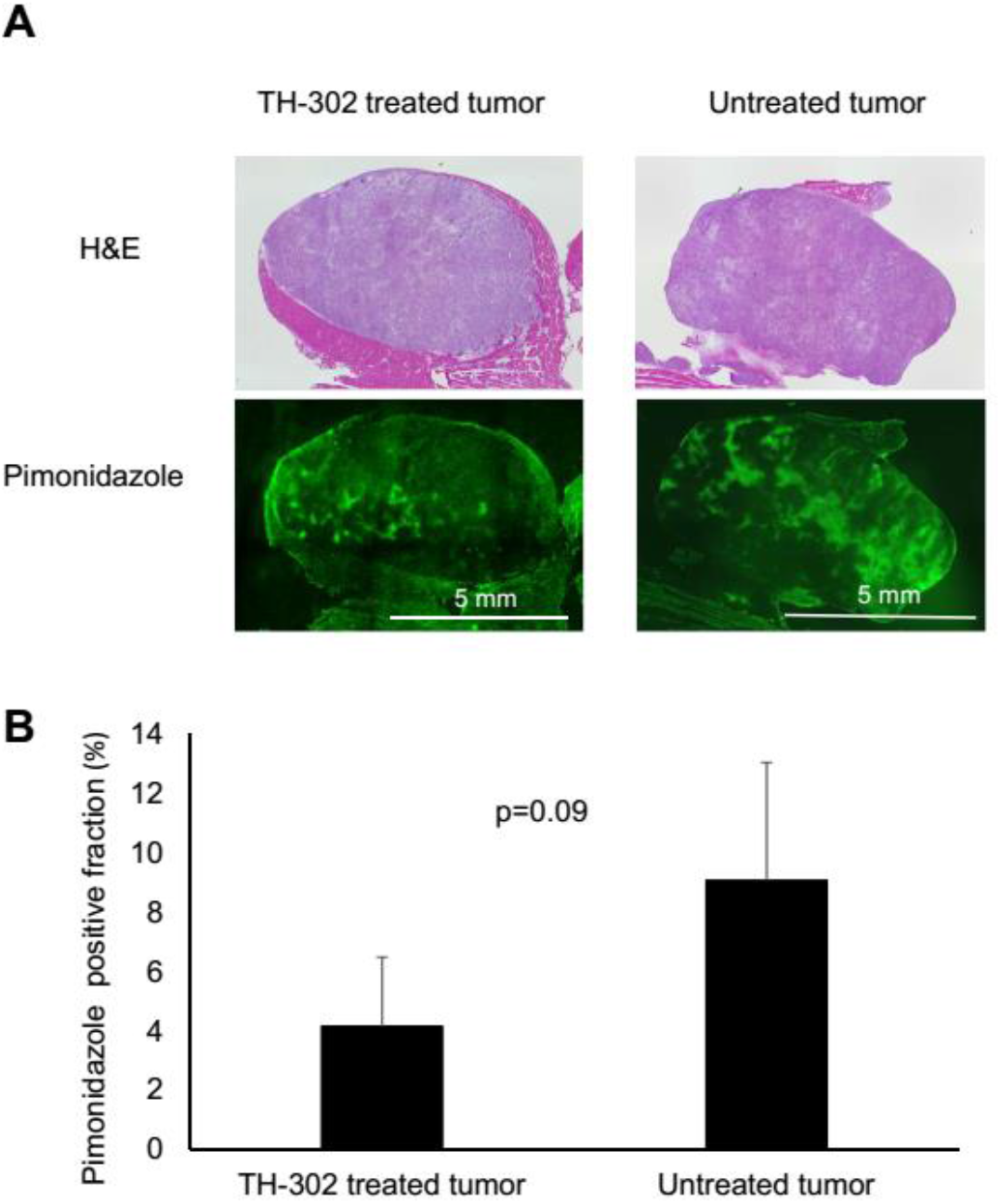
Histological assessment of tumor hypoxia using pimonidazole staining. **A.** representative sets of evofosfamide treated and untreated MIA Paca-2 tumors stained with HE and pimonidazole. **B.** Comparison in pimonidazole positive fraction between evofosfamide treated (n = 4) and untreated MIA Paca-2 tumors (n = 4).

Pimonidazole provides a qualitative measure of hypoxia.^16,21^ To quantify responses to evofosfamide treatment more precisely, we imaged pO_2_ of MIA Paca-2 and Su.86.86 xenografts pre- and post-treatment with evofosfamide by EPR. Figure 5A shows representative images of MIA Paca-2 tumors before treatment (top panels) and after two-day treatment with evofosfamide (bottom panels) while the corresponding frequency histograms of pO2 values before and after treatment for the MIA Paca-2 xenografts for this image are shown in Fig. 5B. No change is evident in any of the tumors for the Su.86.86 evofosfamide insensitive tumor model. The changes for the evofosfamide sensitive MIA Paca-2 tumor is more complex. In some MIA Paca-2 tumors, a reduction in the size of the hypoxic fraction and a shift in the distribution towards higher pO2 levels is evident after treatment. This is especially evident in pre-treatment tumor regions with pO2 in the range 16 and higher. In regions with pO2 <16 mmHg and lower, no significant change or even an increase in hypoxia was detected. The degree to which evofosfamide improves oxygenation is likely correlated with the extent in which the active drug is generated. It is therefore expected that the degree of change should be correlated with the initial level of hypoxia. We therefore plotted the correlation between the initial pO2 and HF10 values and the respective changes in these values. MIA Paca-2 tumors showed a statistically significant negative linear correlation between pre-treatment pO2 and ΔpO2(Δ pO2 = post-treatment pO2 - pre-treatment pO2) (Fig. 5C) and between pretreatment HF10 and ΔHF10 (ΔHF10 = post-treatment HF10 - pre-treatment HF10) (Fig. 5D). The Su.86.86 tumor xenografts by contrast did not show major changes in pO2 and HF10 between untreated and treated tumors (Fig. 5E,4F).

**Figure 5.**
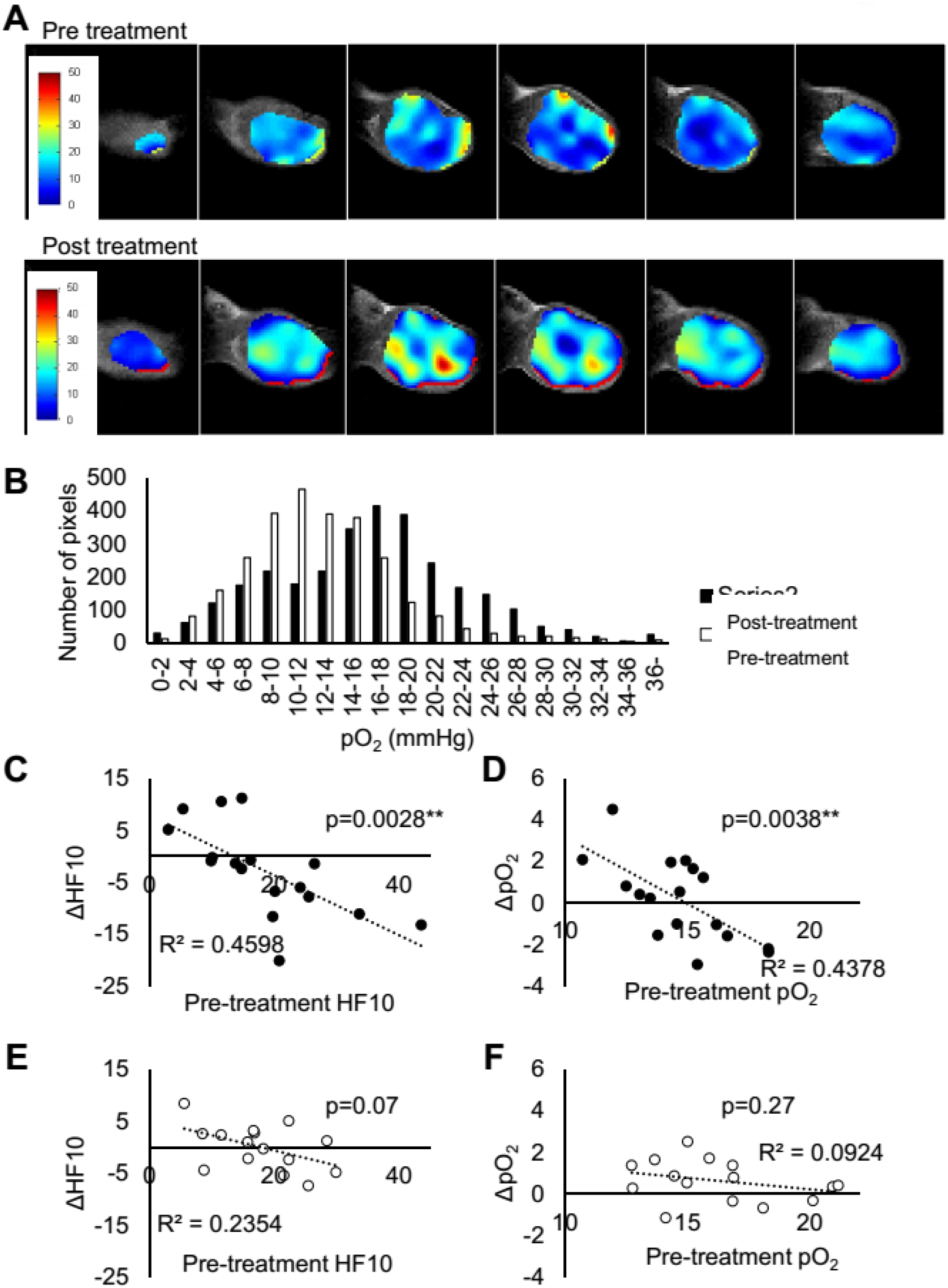
The effect of evofosfamide treatment on EPR oximetry. **A.** representative EPR pO2 images from a MIA Paca-2 tumor taken before and 48 hours after treatment showing the improved oxygenation. **B, C.** the plot of median PO2 before and 48 hours after treatment in MIA Paca-2 tumors and Su.86.86 tumors, respectively. **D, E.** dot plots of pre-treatment HF10 vs ΔHF10 and pre-treatment pO2 vs ΔpO2 in MIP Paca-2 tumors. F, G. dot plots of pre-treatment HF10 vs ΔHF10 and pre-treatment pO2 vs ΔpO2 in Su.86.86 tumors. ** shows p<0.01

Hypoxia can result from either a demand for oxygen that is in excess of a functioning blood supply network or from a defective vascular network delivering an insufficient supply. To determine evofosfamide’s effect on oxygen supply, we investigated the treatment effect of evofosfamide on tumor perfusion and permeability by dynamic contrast enhanced (DCE) MRI using Gd-DTPA as a contrast agent. K^trans^ values were evaluated before treatment and after 2 days of daily treatment of evofosfamide. In the absence of treatment, K^trans^ increased after 2 days in both MIA Paca-2 and Su.86.86 tumors (Fig. 6A, B), and this increase was significantly decreased in both tumor models in the evofosfamide treated mice, consistent with previous reports of decreased K^trans^ by evofosfamide treatment. ^29,30^ We further examined the treatment effect on blood vessels by measuring the CD31 positive fraction in the MIA Paca-2 tumor treated with evofosfamide or vehicle. There was no significant difference in CD31 between evofosfamide and the control group, indicating the vascular damage caused by evofosfamide treatment was minimal (Fig. 6C).

**Figure 6.**
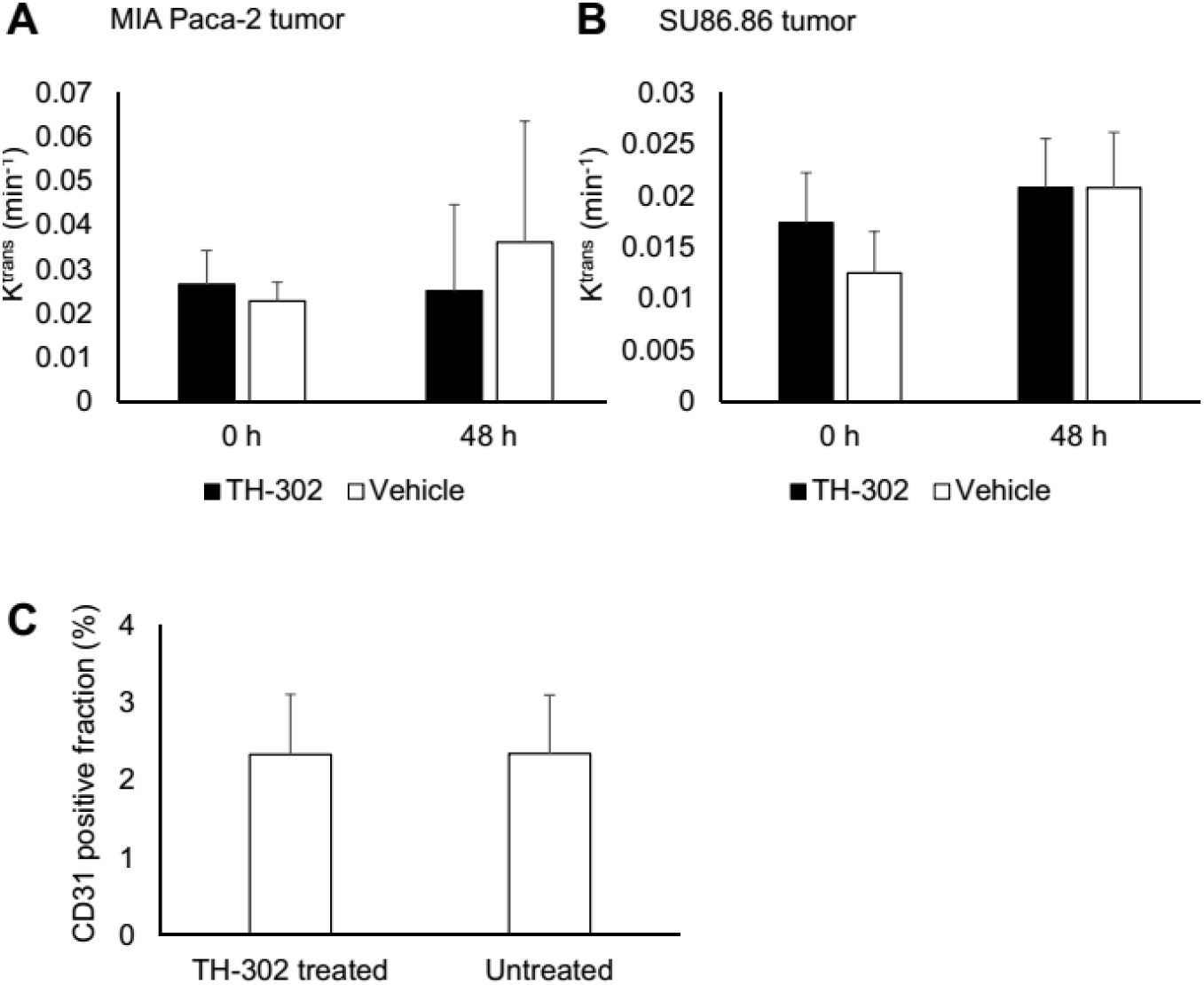
Improved pO2 induced by neither improved perfusion/permeability nor angiogenesis. **A, B** K^trans^ of Gd-DTPA in MIA Paca-2 and Su.86.86 tumors treated with evofosfamide (50 mg/kg daily) or vehicle for 48 hours. **C.** representative of MIA Paca-2 tumor stained with CD 31, αSMA, and DAPI antibodies. **D.** CD 31 positive fraction was measured by histological assessment. Each bar represents four tumor samples.

Evofosfamide is known to have a synergistic effect with radiation therapy (RT) in pancreatic cancers, as evofosfamide targets hypoxic fractions of the tumor and radiation therapy is an oxygen dependent therapy. Both the hypoxic and normoxic fraction of the tumor can be treated in this manner. An additional synergistic effect is possible due to enhanced efficacy of RT in hypoxic regions due to the increase in oxygenation after evofosfamide treatment. A combination treatment of evofosfamide and RT was designed accounting for the increase in tumor oxygenation 2 days after evofosfamide treatment on MIA Paca-2 tumors (Fig. 7A). With this regimen, RT is supposed to be performed after depleting the hypoxic region in the tumor. Figure 7B shows the tumor growth curve of MIA Paca-2 tumors treated with either vehicle, evofosfamide monotreatment, or combination treatment. Although the combination therapy was treated for a shorter period of time, relatively stronger growth inhibition compared to monotherapy was observed in combination therapy group. This result along with EPRI suggests that evofosfamide treatment improves tumor oxygenation, thereby improving the effectiveness of radiotherapy when followed by evofosfamide.

**Figure 7.**
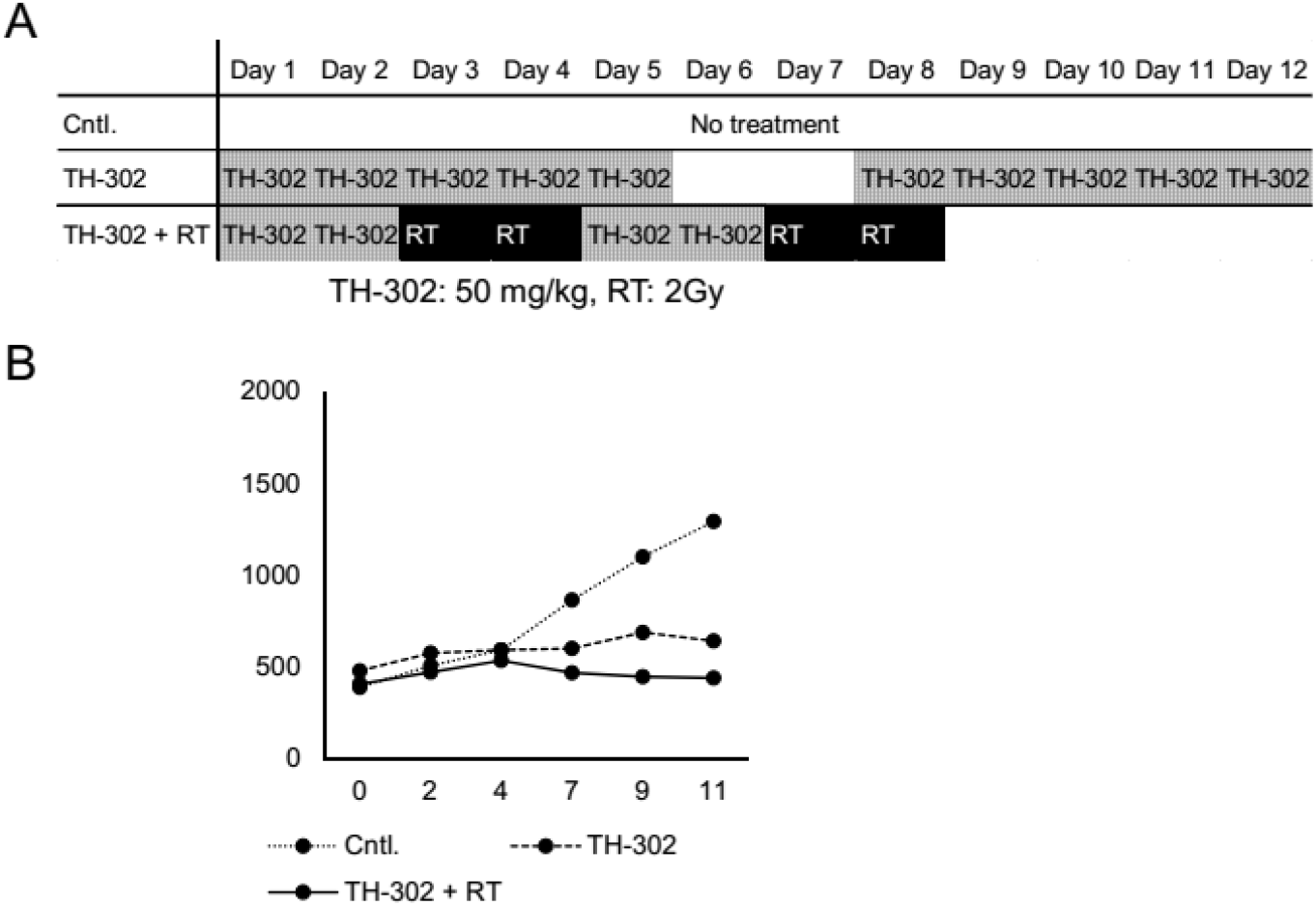
Designing evofosfamide + radiation combination treatment regimen accounting for evofosfamide induced hypoxia. The tumor growth inhibition by evofosfamide + radiation combination treatment regimen accounting for evofosfamide induced oxygenation. **A.** scheme of combination therapy and evofosfamide monotherapy regimens. **B.** the tumor growth curve of no treatment, evofosfamide monotherapy, and evofosfamide +RT combination therapy.

## Discussion

Earlier studies have shown that MIA Paca-2 PDAC tumors are more sensitive to evofosfamide than Su.86.86 tumors. It is believed that the difference is mainly due to the higher hypoxia levels in MIA Paca-2 tumor associated with poor vasculature, since reductive activation of evofosfamide is necessary to release the active fragment of evofosfamide which is thought to occur at cytotoxic levels only in hypoxic regions deep within the tumor. A causative role for hypoxia is suggested by the time course of inhibition, as inhibition of tumor growth is not evident in the first few days (Figs. 2A and 2B) when the small MIA Paca-2 tumors (<600 mm^3^) have pO2 levels comparable to their Su.86.86 counterparts (Fig. 2E).

The correlation with pO2 levels observed is consistent with previous results. However, an underappreciated aspect of evofosfamide treatment is that oxygen levels are not constant during treatment as evofosfamide itself can alter the oxygenation state as apoptosis/necrosis induced by evofosfamide decreases oxygen consumption. Since evofosfamide mechanism is oxygen dependent, this alteration in oxygenation status may potentially decrease the treatment’s effectiveness over time. Despite its potential impact on evofosfamide treatment, only a few studies have evaluated the tumor pO2 after evofosfamide monotherapy, mostly by pimonidazole staining. Most of them showed improved pO2 levels ^8,11,13^ 1-2 day after the treatment while one showed no improvement one day after the treatment.^13^ The study by Peeters et.al employed both pimonidazole staining and [18F] HX4 PET imaging to evaluate the oxygen modification by evofosfamide treatment.^11^ Although both methods qualitatively showed improved oxygenation after evofosfamide treatment, there was a disparity in values between the two modalities probably due to the lack of the capability of quantitative assessment with PET imaging.

EPRI oximetry is a useful imaging modality to clarify this phenomenon because it allows minimally invasive *in vivo* quantitative pO2 assessment. The capability to distinguish quantitatively tumor oxygen status from 5 mm Hg to 25 mm Hg gives this method the capability to assess changes after pharmacologic interventions. By EPRI, we could also observe the effect on pO2 in each unique tumor and identify tumors likely to be susceptible to evofosfamide treatment. In addition to allowing comparison between tumor types, intra-tumor pO2 levels measured by EPRI provide even more detailed information on the effect of evofosfamide treatment on the tumor microenvironment. Not only can the oxygenation effect of evofosfamide therapy be seen in Figure 5B-F, but it can also be seen that this differential effect is dependent on the initial pO2 profile before treatment. The ΔpO2 change pre- and post treatment linearly correlated with pre-treatment pO2 only in MIA Paca-2 tumor (Fig. 5D) and similar relationship was also observed between ΔHF10 and pre-treatment HF10 (Fig. 5C). This is consistent with the capability of evofosfamide to induce cytotoxicity, indicating the causative relationship between evofosfamide treatment and pO2 improvement in tumor. Further, a median pO2 threshold of approximately 15 mmHg, established from the intercept of the trend line on horizontal axis indicates the initial pO2 level at which sensitive-tumors can benefit from evofosfamide treatment (Fig. 5D). It is noteworthy that evofosfamide resistant Su.86.86 tumors didn’t exhibit a pO2 change after treatment even when the pO2 level was comparably low (Fig. 5E, F). This result is consistent with the result of the tumor growth experiment in which evofosfamide was ineffective in Su.86.86 tumors (Fig.2A-D). EPRI is therefore useful in both predicting evofosfamide sensitivity and in monitoring treatment response.

Improvement in oxygenation is often be attributed to improved perfusion. The K^trans^ values of MIA Paca-2 and Su.86.86 xenografts are appreciably different, suggesting higher permeability/perfusion in Su.86.86 xenografts in line with the higher pO2 levels measured by EPRI and in agreement with previous studies.^31^ K^trans^ decreased in both models after treatment, possibly due to reduced VEGF production from selective targeting of the hypoxic fraction.^29^ The decrease in K^trans^ is consistent with a decrease in perfusion/permeability after treatment, which should, other factors be absent, be expected to lead to a decrease in oxygenation. The opposite is in fact observed. This suggests the dominant cause for reoxygenation is not an opening of the vascular network following a decrease in solid state pressure,^32–34^ but rather the direct cytotoxic effect of evofosfamide leading to either cell death or decreased oxygen consumption after cell cycle arrest.

Surprisingly, the relative sensitivity observed *in vivo* was also mirrored *in vitro* under aerobic conditions. Using a long drug exposure model (48 hours, comparable to continuous i.p. administration regimen in vivo) in air-equilibrated plates we evaluated hypoxia-independent toxicity *in vitro* under highly aerobic conditions (Fig. 3). Compared to the control group, MIA Paca-2 cells were still strongly inhibited relative to Su.86.86 cells at all the concentrations examined. Since both cell lines are under ambient air conditions in this experiment, this finding suggests an additional cellular mechanism of treatment resistance independent of oxygen delivery exists in Su.86.86. Furthermore, Su.86.86 cells have an oxygen consumption rate roughly twice that of Mia Paca-2 cells.^14,35^ On the basis of oxygen consumption alone, Su86.86 would be expected to be more sensitive to evofosfamide than Mia Paca-2, the opposite of what is observed. An oxygen independent difference must therefore exist. Many possibilities exist including differences in the activity of one-electron reductases,^36^ differences in the efficiency of DNA repair,^7,14^ and differences in cell fate as a function of DNA damage. These differences in cellular metabolism are not readily accessible by imaging and must be accounted for by other methods.

Several regimens for evofosfamide combination therapy have been examined in preclinical studies and proven to be effective.^10,12,19^ Such preclinical studies usually used a frequent dosing of evofosfamide. Based on these studies, clinical trials also used weekly dosing regimens. As a result of such frequent dosing, gastrointestinal disorders such as nausea and vomiting were commonly observed in the phase I clinical trials for evofosfamide, ^9^ which may limit compliance. As the decreased hypoxic fraction following evofosfamide treatment sensitizes the tumor to radiation, the synergistic action may reduce the needed dose of evofosfamide. This hypothesis was examined in the experiment comparing the combination regimen with extended monotherapy regimen in Figure 6A. The result showed that alternating treatment of the combination therapy showed a comparative effect to extended monotherapy with evofosfamide (Fig. 7B). This suggested that when planning the combination therapy of radiation and evofosfamide, decreasing frequency of evofosfamide treatment may decrease the systemic toxicity without compromising the treatment effect. With methodologies to provide quantitative assessment of tumor oxygen with T1-weighted MRI on conventional clinical scanners implemented on human subjects ^37,38^, such strategies will play a key role in tailoring therapies with evofosfamide and in combination with antiproliferative therapies such as conventional chemotherapy or radiation therapy.

## Conclusions

This study provides strong evidence of decreasing hypoxic fraction exclusively from hypoxic tumor. The result from EPRI oximetry suggested the importance of evaluating hypoxia in individual tumors. This hypoxia decreasing effect may be utilized in designing regimens for the combination therapy of evofosfamide and radiation therapy. MRI methods to provide quantitative assessment of tumor oxygenation and changes in response to treatment will play a key role in combination treatments.

